# *N*^6^-methyladenosine and the NEXT complex direct Xist RNA turnover and X inactivation dynamics

**DOI:** 10.1101/2025.01.09.632176

**Authors:** Guifeng Wei, Heather Coker, Lisa Rodermund, Mafalda Almeida, Holly L. Roach, Tatyana B. Nesterova, Neil Brockdorff

## Abstract

X chromosome inactivation (XCI) in mammals is orchestrated by the non-coding RNA Xist which together with specific interacting proteins, functions *in cis* to silence an entire X chromosome. Defined sites on Xist RNA carry the *N*^6^-methyladenosine (m^6^A) modification, and perturbation of the m^6^A writer complex has been found to abrogate Xist-mediated gene-silencing. However, the relative contribution of m^6^A and its mechanism of action remain unclear. Here we investigate the role of m^6^A in XCI by applying rapid degron-mediated depletion of METTL3, the catalytic subunit of the m^6^A writer complex, an approach that minimises indirect effects due to transcriptome-wide depletion of m^6^A. We find that acute loss of METTL3/m^6^A accelerates Xist-mediated gene silencing, and that this correlates with increased levels and stability of Xist transcripts. We show that Xist RNA turnover is mediated by the nuclear exosome targeting (NEXT) complex but is independent of the principal nuclear m^6^A reader protein YTHDC1. Our findings demonstrate that the primary function of m^6^A on Xist RNA is to promote Xist RNA turnover which in turn regulates XCI dynamics.

## Introduction

X chromosome inactivation (XCI) is a developmentally regulated process that evolved in mammals to equalise the levels of X-linked gene expression in XX females relative to XY males^1^. Silencing of a single X chromosome in cells of XX embryos is orchestrated by the X inactive specific transcript (Xist), an ∼17 kb non-coding RNA, which accumulates *in cis* across the future inactive X (Xi) chromosome^2–5^. Functional elements within Xist RNA have been assigned in large part to tandem repeat blocks labelled A-F, that are distributed across the length of the transcript. Most notably, the 5’-end located A-repeat element has been found to be critical for Xist-mediated gene silencing^6^. Identification of RNA binding proteins (RBPs) that interact with the A-repeat and other elements has been achieved using both proteomic^7–9^ and functional genetic screening^10,11^ strategies. These approaches led to the identification of the RBP SPEN as a critical factor for Xist-mediated silencing, functioning via binding to Xist A-repeat region^12^ and recruitment of the NCoR-HDAC3 histone deacetylase complex^7,13^. The Polycomb system, which contributes to Xi silencing, is recruited via the RBP hnRNPK which binds to the Xist B/C-repeat region^14,15^. Another RBP, the SPEN related protein RBM15, was identified both in proteomic and functional screening^8,10^. Follow-up studies have shown that RBM15 is an accessory subunit of the multiprotein complex that catalyses RNA *N*^6^-methyladenosine (m^6^A)^16^. Other subunits of the m^6^A writer complex were also identified in the Xist proteomic and functional screening experiments^8,10^.

Building on initial observations implicating the m^6^A writer complex in XCI, it was reported that RBM15 directs m^6^A activity to two sites immediately downstream of the Xist A-repeat and E-repeat regions, and that perturbation of the complex, or of the protein YTHDC1, which binds to m^6^A modified RNA in the nucleus, strongly abrogates Xist-mediated gene silencing^16^. In contrast, other studies that analysed Xist-mediated gene silencing reported minor or negligible effects on XCI following knockout of genes encoding subunits of the m^6^A writer complex^17^ or following deletion of m^6^A sites in the vicinity of the Xist A-repeat^17,18^. Likely explanations for these discrepancies include the use of different cell models, different assays to assess Xist-mediated silencing and different strategies for perturbation of the m^6^A writer complex^19^. Of particular note is that m^6^A impacts the function of several thousand mRNAs such that chronic or long-term gene knockout studies have the potential to lead to significant secondary or indirect effects.

In this study we exploit an alternative approach, acute protein depletion with the dTAG degron system^20^, to investigate the role of the m^6^A writer complex in Xist-mediated silencing. Our study demonstrates that the primary function of m^6^A on Xist RNA is to promote transcript turnover and that removal of METTL3 increases the rate of Xist-mediated silencing. Additionally, we find that the NEXT (nuclear exosome targeting) complex is essential for Xist degradation via a pathway that functions independently of the major nuclear m^6^A reader protein YTHDC1.

## Results

### Acute depletion of METTL3 accelerates Xist-mediated gene silencing

In recent work we analysed the role of m^6^A in regulating nascent RNA processing, making use of the dTAG degron system^21^ to rapidly deplete METTL3, the catalytic subunit of the m^6^A writer complex, in iXist-ChrX_Cast_ (clone C7H8) mouse embryonic stem cells (mESCs)^17^. We showed that METTL3 depletion results in rapid loss of m^6^A across the transcriptome, including over Xist m^6^A peak regions in proximity to the A-repeat and E-repeat^21^. Here, we have employed the same degron strategy to investigate how acute loss of METTL3/m^6^A affects Xist-mediated chromosome silencing. iXist-ChrX_Cast_ mESCs have an interspecific *Mus castaneus* x 129 strain background, a stable XX karyotype and have been engineered with a TetOn promoter for induction of Xist expression specifically from the *Mus castaneus* X chromosome^17^. Accordingly, we are able to isolate chromatin-associated RNA and subject it to sequencing (ChrRNA-seq) to accurately determine the relative expression level of active X (Xa) and inactive X (Xi) alleles for a large number of X-linked genes that have informative SNPs. In addition to two previously described lines with a METTL3 C-terminal FKBP12^F36V^ tag^21^, we derived two independent lines with METTL3 tagged with FKBP12^F36V^ on the N-terminus (Fig. 1a) in the iXist-ChrX_Cast_ background. In all FKBP12^F36V^ tagged cell lines METTL3 depletion occurred rapidly, within 2 hours of adding the dTAG-13 reagent (Fig. 1b and Extended Data Fig. 1a), and moreover resulted in strongly reduced levels of METTL14, a subunit of the m^6^A complex that heterodimerises with METTL3(Fig. 1b and Extended Data Fig. 1b). We noted a reduction in levels of N-terminal METTL3-FKBP12^F36V^, indicating that insertion of the tag affects METTL3 protein stability (Extended Data Fig. 1c).

**Figure 1:**
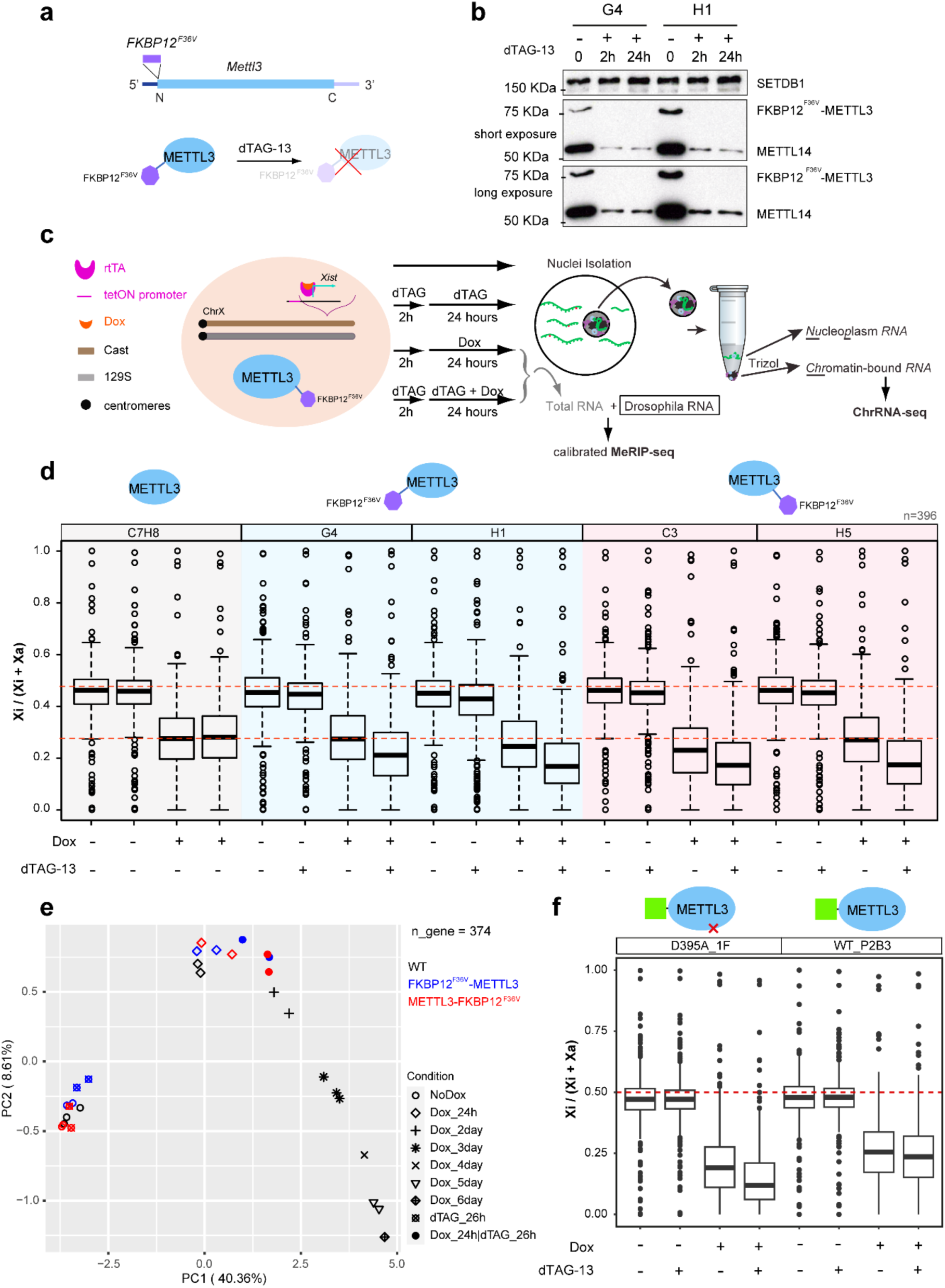
Acute depletion of METTL3 results in accelerated X chromosome inactivation. **a**, Schematic outline of N-terminal FKBP12^F36V^ tagging of METTL3. **b**, Western blot showing the FKBP12^F36V^-METTL3 fusion protein and METTL14 protein level for two independent clones (G4 and H1) after 2 or 24 h dTAG-13 treatment. SETDB1 loading control. Middle and lower panel represent short and long exposure respectively. **c,** Schematic outline detailing cell line specification and experimental design. **d**, Boxplot showing the allelic ratio of X-linked genes (n=396) from ChrRNA-seq analysis for each sample and condition indicated above and below respectively. Two independent N-terminal degron (G4 and H1 clone) and C-terminal FKBP12^F36V^ (C3 and H5 clone) tagged lines are included for this analysis, alongside untagged wild-type (WT) cells (C7H8 clone). The Y-axis denotes allelic ratio ranging from 0 to 1. Two red dashed lines indicate allelic ratio in ES cells (NoDox) and one day dox-induced Xist condition for WT cells. **e**, PCA using allelic ratio of X-linked genes (n=374) for samples in panel b, along with time-course WT (C7H8) samples. **f**, Boxplot depicting the allelic ratio of X-linked genes from ChrRNA-seq analysis for the complementation assay where either GFP-METTL3 (WT_P2B3 clone) or GFP-METTL3^D395A^ (D395A_1F clone) is expressed from the *Rosa26* locus in a C-terminal METTL3 dTAG degron cell line (H5). Both WT_P2B3 and D395A_1F clone retain both X_cast_ and X_129_. The red dashed line indicates allelic ratio at 0.5. Samples and conditions are indicated above and below respectively.

We went on to quantify Xist-mediated silencing following METTL3 depletion as outlined in Fig. 1c. We analysed silencing at an early time-point, 24 h Xist induction (with or without prior dTAG-13 treatment for 2 h), in order to minimise potential indirect effects of m^6^A depletion (Xist-mediated silencing in the iXist-ChrX_Cast_ mESCs occurs progressively over a period of around 6 days^22^). The results are summarised in Fig. 1d. In the absence of dTAG-13 reagent, silencing levels in FKBP12^F36V^ tagged lines were very similar to those seen in control C7H8 cells, indicating that there are no effects attributable to addition of the degron at either the C- or N-terminus of METTL3.

Interestingly, addition of dTAG-13 resulted in a highly reproducible enhancement or acceleration of Xist-mediated silencing, evident in all four independently derived clones (Fig. 1d). Of note, accelerated silencing is the converse of the silencing deficiency reported in prior work using constitutive knockout or knockdown of METTL3 and of other subunits of the m^6^A writer complex^16,17^. This difference likely reflects that acute METTL3 depletion is less influenced by indirect effects compared to chronic knockout experiments. Indeed, global gene expression differences in our study correlate better with m^6^A modified mRNAs compared to published studies that used long-term knockout approaches^23^ (Extended Data Fig. 1d). Principal component analysis comparing silencing in untreated cells with acute depletion of METTL3 after 24 h Xist induction (Fig. 1e), together with silencing analysis for previously defined gene categories^15,17,22^ (Extended Data Fig. 1e-h), indicate that all X-linked genes are equally subject to accelerated silencing.

To determine if the observed acceleration of Xist-mediated silencing is attributable to loss of METTL3 catalytic function, we performed complementation experiments using ectopic expression of GFP-METTL3 transgenes. Transgene constructs were integrated under the control of the Rosa26 constitutive promoter into the H5 cell line with C-terminal degron tagged METTL3 using CRISPR/Cas9 facilitated homologous recombination. In parallel, we established lines using a transgene encoding GFP-METTL3^D395A^, a point mutation that ablates METTL3 catalytic activity^24^. Constitutive expression of GFP-METTL3 transgenes was maintained both in the presence and absence of dTAG-13 (Extended Data Fig. 2a-c).

Both transgene encoded proteins reversed the observed reduction in METTL14 protein levels (compare Extended Data Fig. 2b,c with Fig. 1b and Extended Data Fig. 1b), indicating the formation of stable GFP-METTL3/METTL14 heterodimers. Compared to wild-type (WT) GFP-METTL3, levels of GFP-METTL3^D395A^ were reduced (Extended Data Fig. 1c, Fig.2b,c). This effect was not seen in the presence of dTAG-13 and is not linked to transcriptional levels (Extended Data Fig. 2d). A possible explanation is that METTL3 dependent m^6^A autoregulates the cDNA-derived ectopic *Mettl3* transcript via RNA degradation. We went on to analyse Xist-mediated silencing in the transgene complementation lines. ChrRNA-seq analysis showed that ectopic expression of WT GFP-METTL3 fully complements the accelerated silencing phenotype observed following dTAG-13 treatment, whilst expression of GFP-METTL3^D395A^ had no effect (Fig. 1f, Extended Data Fig. 2e). Collectively, these results confirm the role of METTL3 in regulating the rate of Xist-mediated silencing and further indicates that METTL3 function in this circumstance is via catalysis of m^6^A.

### Accelerated XCI is linked to increased levels and stability of Xist RNA

We noted a close correlation between accelerated Xist-mediated silencing and levels of Xist RNA, as determined from ChrRNA-seq datasets. Specifically, increased Xist RNA levels up to approximately 2-fold were apparent following acute METTL3 depletion in the independent N-terminal and C-terminal tagged cell lines (Fig. 2a). Additionally, complementation with WT GFP-METTL3 but not GFP-METTL3^D395A^ restored Xist RNA levels to those seen in untreated cells (Fig. 2b and Extended Data Fig. 2f).

**Figure 2:**
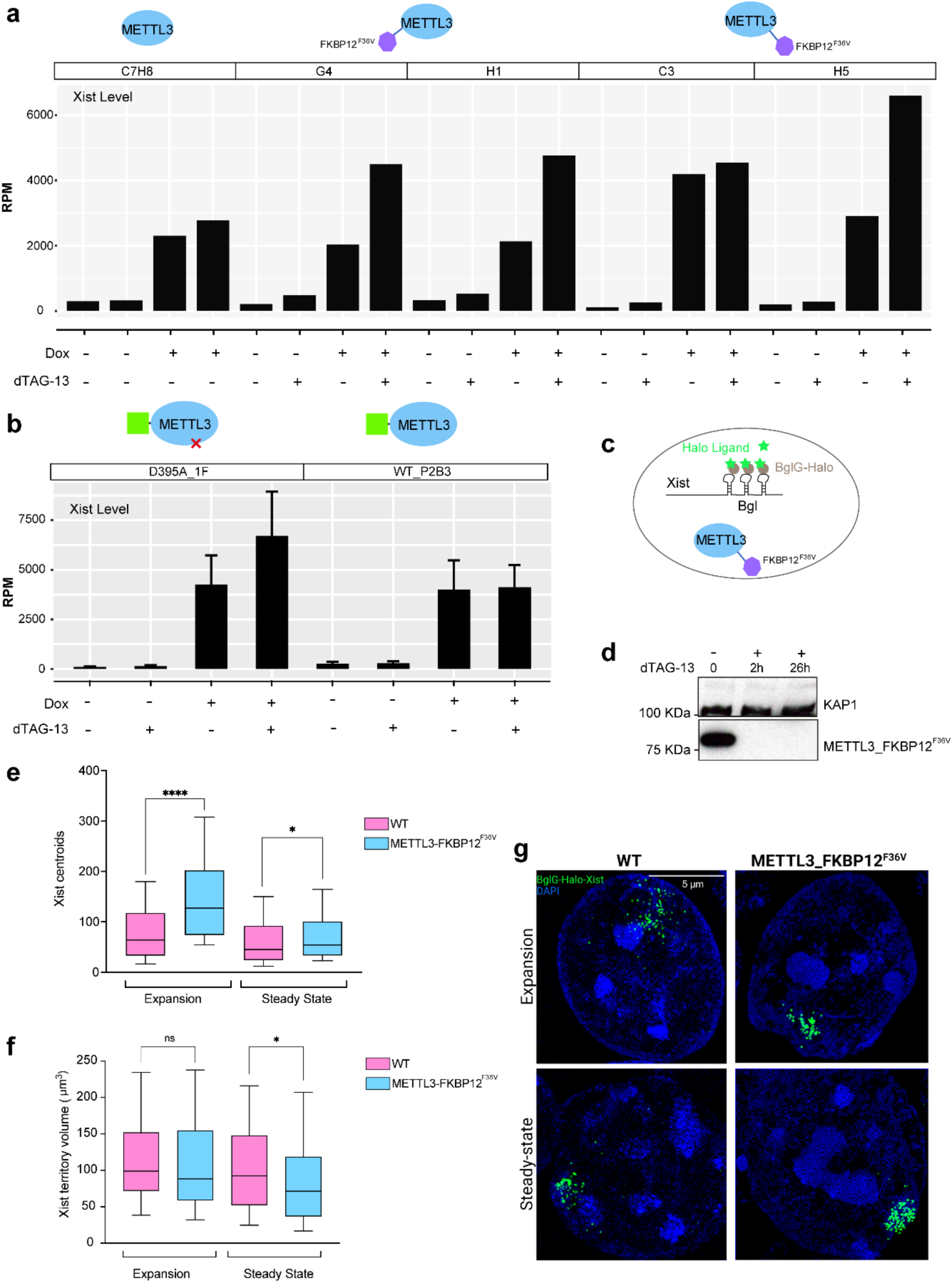
Xist RNA levels are elevated following acute depletion of METTL3. **a**, Barplot showing the expression level (RPM, reads per million mapped reads) of Xist from ChrRNA-seq analysis for each sample and condition described in Fig. 1d. **b**, Expression level of Xist, as in (a), from ChrRNA-seq analysis for samples and conditions described in Fig. 1f. **c**, Schematic of cell line with METTL3 dTAG degron and Xist exon 7 18x Bgl stem-loop bound by BglG-Halo fusion protein for fluorescent imaging of Xist RNA. **d**, Western blot showing rapid degradation of METTL3-FKBP12^F36V^ upon dTAG-13 treatment in the cell line depicted in (c). KAP1 is the loading control. **e,f,** Boxplots showing the number of Xist molecules (e) and Xist cloud volume (f) analysed by RNA-SPLIT for both WT, (pink, data from ref^25^) and METTL3 dTAG depleted cells (blue) at both expansion phase (1.5 h Xist induction) and steady-state phase (24 h Xist induction), with n = a minimum of 123 cells. Significance was determined by use of a two-tailed Mann Whitney test (ns = not significant, * p= 0.0135, **** p= <0.0001). **g,** Representative 3D-SIM images (maximum projection) of Xist RNA (HaloTag, green) in WT (data from ref^25^) and METTL3-FKBP12^F36V^ cells at both expansion and steady-state phases. DNA is counterstained with DAPI (blue).

To further investigate the effect of acute METTL3 depletion on Xist RNA we used super-resolution three-dimensional structured illumination microscopy (3D-SIM) imaging to assay features of Xist RNA domains at the single cell level. For these experiments we engineered the METTL3 C-terminal degron into previously described iXist-ChrX_129_ XX mESCs in which the TetOn inducible Xist allele has a Bgl-stem loop array integrated into Xist exon 7, allowing detection of Xist RNA molecules via binding of a BglG-HaloTag fusion protein labelled with fluorescent Halo dyes^25^ (Fig. 2c,d). Using this approach, we observed an increase in the number of Xist molecules following acute METTL3 depletion (Fig. 2e), in agreement with the ChrRNA-seq data. The volume encompassing Xist centroids (overall Xist domain size) was broadly unchanged (expansion phase, 1.5 h Xist induction) or minimally reduced (steady-state phase, 24 h Xist induction) (Fig. 2f). These observations confirm increased levels of Xist RNA and suggest that Xist domains corresponding to the Xi territory remain broadly unchanged.

We also investigated if other well characterised nuclear lncRNAs that are m^6^A modified are similarly affected by acute METTL3 depletion. Accordingly we examined levels of Neat1, Malat1, and Kcnq1ot1 RNAs, all of which are expressed in mESCs and have high levels of METTL3-dependent m^6^A (Extended Data Fig. 3a-c). Levels of Neat1 and Malat1 were unaffected by acute METTL3 depletion, but similar to Xist RNA, Kcnq1ot1 levels increased approximately 1.5-2-fold (Extended Data Fig. 3d-f). The effect on Kcnq1ot1 levels was dependent on METTL3 catalytic activity (Extended Data Fig. 3f, right).

Increased abundance of Xist RNA following acute depletion of METTL3 could result from changes in the rate of Xist RNA transcription and/or RNA turnover. To investigate these possibilities, we applied RNA-SPLIT (sequential pulse localization imaging over time) coupled to super-resolution 3D-SIM microscopy^25^ to differentially label successive waves of Xist transcripts (pre-synthesized and newly synthesized) prior to fixation. The labelling regimen for determining turnover rates is shown in Fig. 3a. Experiments were performed at both expansion phase and steady-state phase using a 20-minute interval. Two-hour dTAG-13 treatment was performed before Xist induction to minimise secondary/indirect effects. Example images in Fig.3b are from expansion phase. As shown previously, turnover of Xist RNA occurs within 140 min in expansion phase and 220 min at steady-state phase^25^ (Fig. 3b,c). In marked contrast, following acute METTL3 depletion there was little Xist RNA turnover detectable across the entire time course of the experiment (220 min), both during expansion and steady-state phases (Fig. 3b,c). Reduced turnover of Xist transcripts was also demonstrated using an orthogonal approach, SLAM-seq^26^, based on transient 4sU labelling of newly synthesised RNA (Extended Data Fig. 4a-c). Allele-specific analysis using these sequencing data confirmed accelerated silencing and increased Xist RNA levels following acute METTL3 depletion (Extended Data Fig. 5).

**Figure 3:**
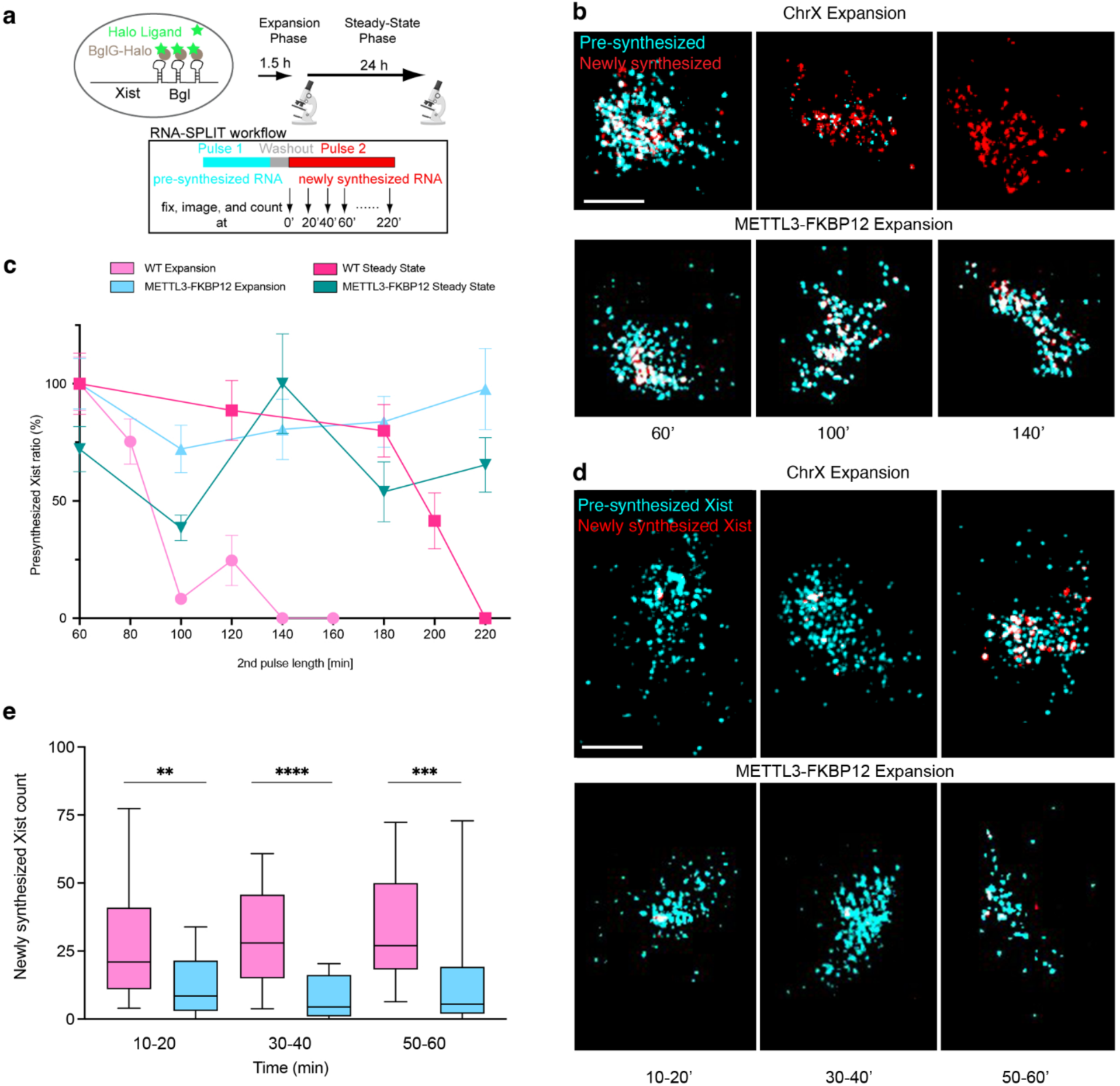
Acute depletion of METTL3 stabilises Xist RNA. **a**, Schematic detailing RNA-SPLIT to assess the rate of Xist RNA turnover. **b**, Representative 3D-SIM images illustrating z-project of pre-synthesized (cyan) and newly synthesized (red) Xist molecules during expansion phase for WT (data from ref^25^) and METTL3 dTAG cells. Images representative of 60, 100, and 140 minutes of pulse 2 labelling, scale bar; 2 μm. **c**, Plot showing quantification of Xist RNA turnover during expansion and steady state for WT (pink, data from ref^25^), METTL3 dTAG (blue), *n* = minimum of 20 cells per time point. **d**, As in (b) but images representative of pulse 2 labelling at indicated times, scale bar; 2 μm. **e**, Boxplot showing Xist RNA transcription over time during expansion phase for WT (pink, data from ref^25^) and dTAG-13 treated METTL3 degron cells (blue). *n* = minimum of 22 cells per time point. Significance was determined by use of a two-tailed Mann Whitney test (** p= 0.0091, *** p=0.008, **** p= <0.0001).

We further applied RNA-SPLIT to measure Xist transcription rates in the presence and absence of m^6^A, achieved by quantifying foci for newly synthesized Xist RNA over time during expansion phase. As shown in Fig. 3d,e, there is a significantly reduced transcription rate in the acute METTL3 depletion condition compared to WT cells. This finding is consistent with prior work demonstrating a feedback mechanism that links Xist transcription and turnover^25^. Accordingly, we conclude that loss of m^6^A results in accelerated X chromosome silencing due to over-accumulation of Xist transcripts.

### Xist RNA turnover is mediated by the ZCCHC8/NEXT complex independent of YTHDC1

The cellular functions of m^6^A are mediated by reader proteins that can bridge to downstream pathways. YTHDC1 is the best characterized protein that directly recognises m^6^A in the nucleus^27^. Interestingly, YTHDC1 co-immunoprecipitation experiments have revealed an association with ZCCHC8, a core subunit of the the nuclear exosome targeting complex (NEXT) complex that targets non-polyadenylated transcripts in the nucleus for degradation^28^. Both YTHDC1 and ZCCHC8 play a role in degrading non-polyadenylated chromatin-associated regulatory (car) RNAs, for example PROMPTs and eRNAs^29,30^. Consistent with this finding, ZCCHC8 has been reported to interact with YTHDC1 in experiments using stable isotope labelling in cell culture (SILAC) and mass spectrometry^28^. The YTHDC1-RNA exosome axis has also been implicated in the degradation of other nuclear RNAs, for example immunoglobulin heavy chain locus-associated lncRNA (SμGLT)^31^ and *C9ORF72* repeat RNA^32^.

To investigate if YTHDC1 is important for regulating Xist RNA turnover, we used CRISPR-Cas9 facilitated genome editing to establish XX mESC derived lines with the FKBP12^F36V^ degron tag inserted into the gene encoding YTHDC1 (Fig. 4a). YTHDC1 depletion on addition of dTAG-13 reagent was validated by western blot analysis (Fig. 4b,c) and also by examination of the effects on *Tor1aip2* alternative last exon splicing which is significantly affected by METTL3/m^6^A and YTHDC1 depletion^21^ (Extended Data Fig. 6a,b). We went on to assay Xist-mediated silencing and Xist RNA levels following YTHDC1 depletion as described above. We observed no increase in the silencing rate nor in levels of Xist RNA in two independent degron tagged cell lines (Y9F and Y12H) (Fig. 4d,e), both if anything being slightly reduced compared to controls. There was little or no effect on Xist RNA stability as determined by SLAM-seq (Extended Data Fig. 4c). Similarly, Kcnq1ot1 RNA, levels of which increase following METTL3 depletion, were unaffected by YTHDC1 depletion (Extended Data Fig. 3g-i, left).

**Figure 4:**
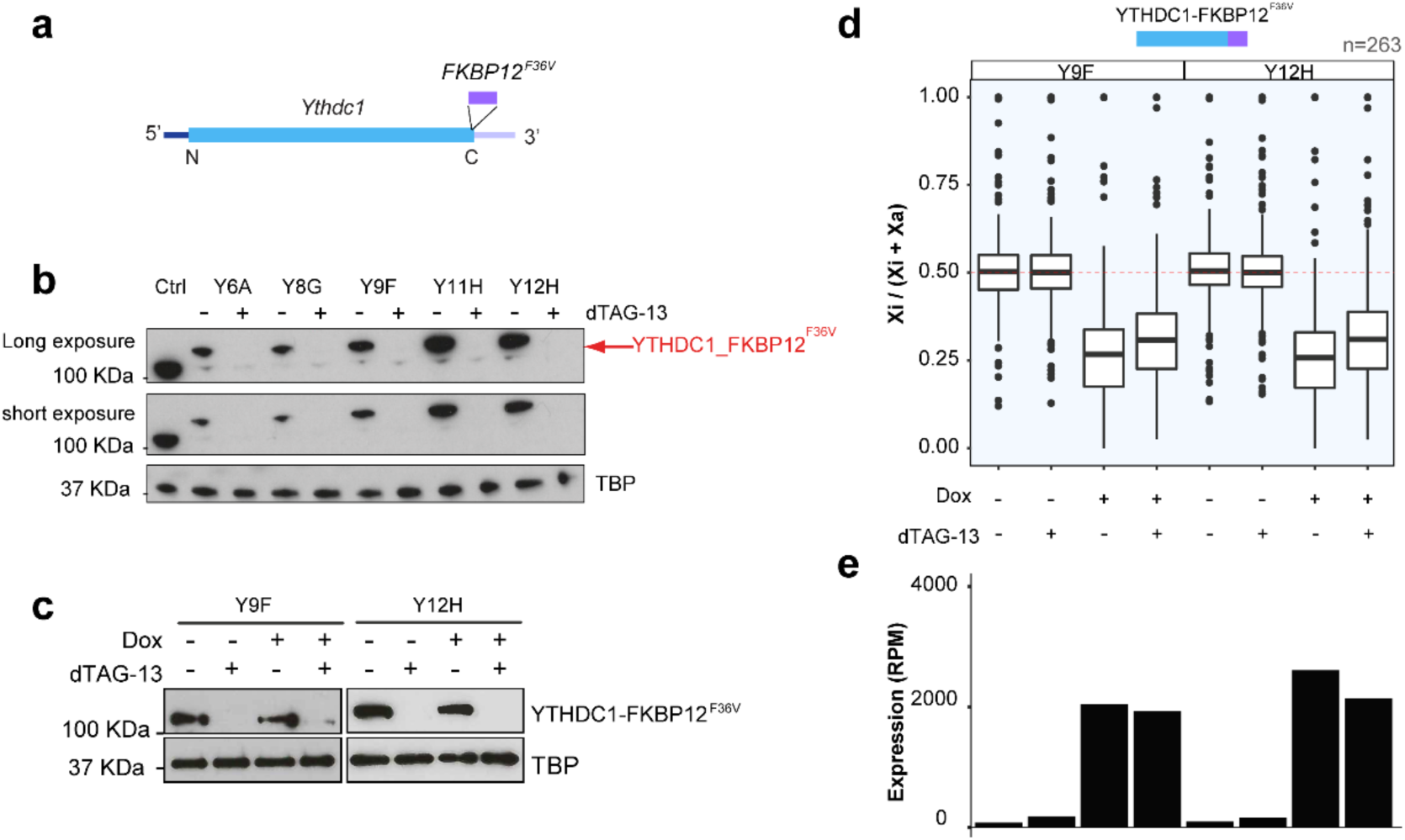
Xist RNA turnover is independent of the m^6^A-nuclear reader protein YTHDC1. **a**, Strategy for in-frame insertion of FKBP12^F36V^ into YTHDC1. **b**, Western blot showing the protein size and level of YTHDC1 upon 0 or 2 h dTAG-13 treatment. The tagged protein is indicated by the red arrow. TBP loading control. **c**, Western blot showing the protein level of YTHDC1-FKBP12^F36V^ for ChrRNA-seq samples shown in Fig. 4d. TBP loading control. **d**, Boxplot showing the allelic ratio of X-linked genes (n=263) from ChrRNA-seq analysis for YTHDC1 dTAG degron samples. The experimental design is as described in Fig. 1a. The red dashed line indicates allelic ratio at 0.5. Samples (two independent clones, Y9F and Y12H) and conditions are indicated above and below respectively. **e,** Barplot showing the expression level of Xist from ChrRNA-seq analysis for samples and conditions described in (d).

We went on to investigate the role of the NEXT complex in Xist RNA turnover by inserting the FKBP12^F36V^ degron tag into the gene encoding the core subunit ZCCHC8 in XX mESCs (Fig. 5a and Extended Data Fig. 7a,b). As an additional control we established XX mESC lines with the FKBP12^F36V^ degron tag inserted into the gene encoding ZFC3H1, an essential subunit of Poly(A) tail exosome targeting complex (PAXT)^33^ (Fig. 5b and Extended Data Fig. 7a,c). PAXT mediates an alternate pathway for degradation of polyadenylated RNA in the nucleus, potentially functioning as a timer to remove aberrant RNAs that are not efficiently exported^33^. dTAG-13 treatment led to rapid and complete depletion of the FKBP12^F36V^ tagged proteins within 2 hours (Fig. 5a,b). We noted that m^6^A-dependent alternative last exon splicing of the *Tor1aip2* gene was not affected by acute depletion of ZCCHC8 or ZFC3H1, indicating that neither NEXT nor PAXT is required for m^6^A deposition on target mRNAs (Extended Data Fig. 7d,e). Acute ZCCHC8 depletion was further validated by monitoring upregulation of PROMPTs (Extended Data Fig. 8a,b).

**Figure 5:**
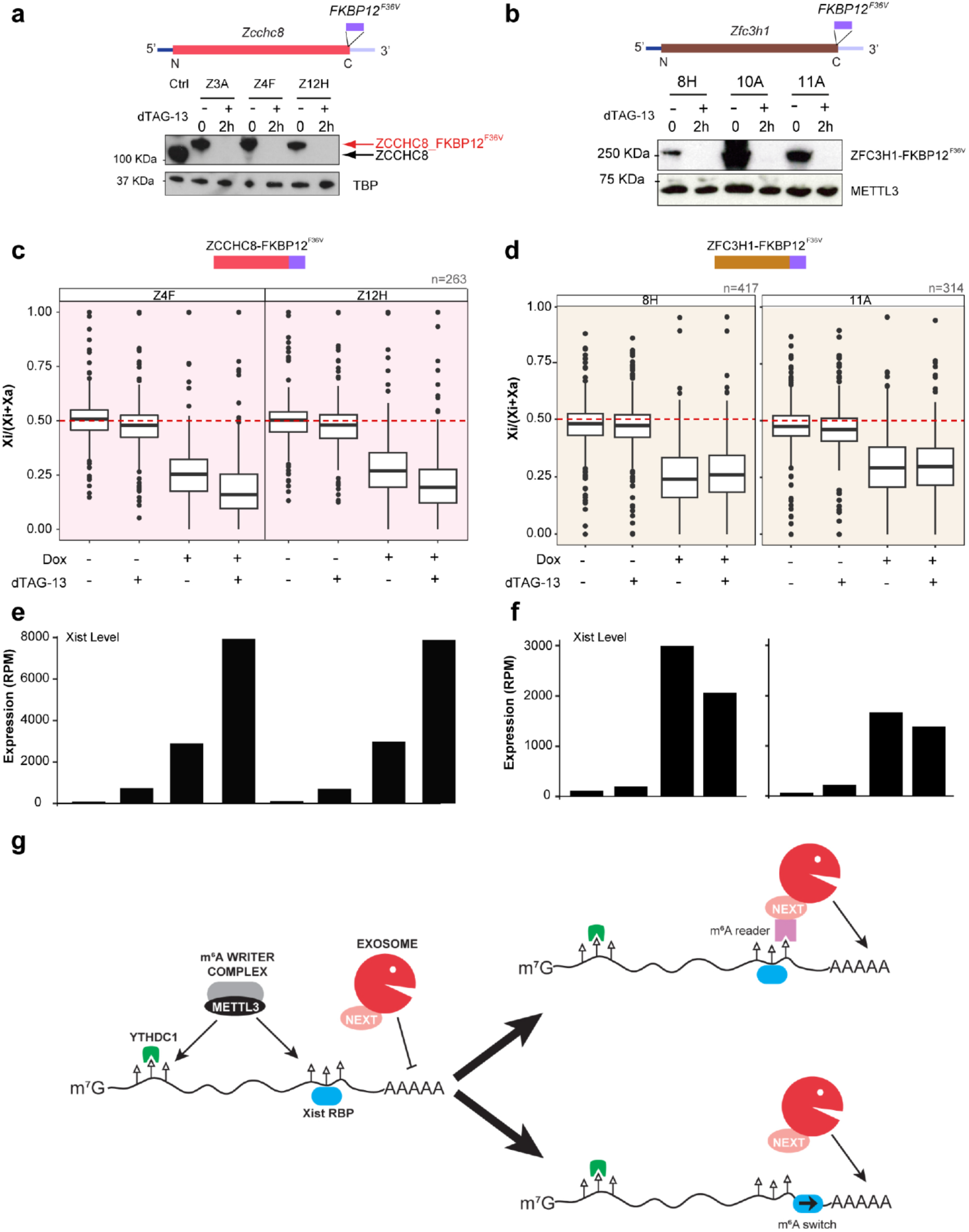
The NEXT complex mediates Xist RNA turnover in the cell nucleus. **a**, Strategy showing in-frame insertion of FKBP12^F36V^ into ZCCHC8 (top). Western blot (bottom) shows the protein level of ZCCHC8-FKBP12^F36V^ after 0 or 2 hours dTAG-13 treatment. TBP loading control. The red and black arrows indicate the protein size for ZCCHC8-FKBP12^F36V^ and untagged ZCCHC8. **b**, As in (a), but for ZFC3H1. METTL3 loading control. **c**, Boxplot showing the allelic ratio of X-linked genes from ChrRNA-seq analysis for ZCCHC8 dTAG degron samples. The experimental design is as described in Fig. 1c. The red dashed line indicates allelic ratio at 0.5. Samples (two independent clones, Z4F and Z12H) and conditions are indicated above and below respectively. **d**, As in (c), but for ZFC3H1. Note that only the proximal 138 MB of X chromosome in 11A clone is informative for allelic-speific analysis. **e,** Barplot showing the expression level of Xist from ChrRNA-seq analysis for samples and conditions described in (c). **f,** As in (e), but for ZFC3H1 clones described in (d). **g,** Schematic depicting alternative models for m^6^A-mediated regulation of Xist RNA turnover as discussed in the main text. m^6^A sites are indicated as lollipops with open triangle.

We went on to perform ChrRNA-seq to assay Xist-mediated silencing after 24 h of Xist induction, either in the presence or absence of NEXT or PAXT complexes using the approach described above for analysis of METTL3 and YTHDC1. As shown in Fig. 5c, acute depletion of ZCCHC8 resulted in strongly accelerated silencing, whereas depletion of ZFC3H1 resulted in a marginal reduction in gene silencing (Fig. 5d). Consistent with these observations, Xist RNA levels were elevated following depletion of ZCCHC8 but not of ZFC3H1, where if anything Xist levels were slightly lower (consistent with marginally reduced gene silencing) (Fig. 5e,f). The higher Xist RNA levels and stability following acute ZCCHC8 depletion is correlated with an even more marked acceleration of silencing that was seen with acute METTL3 depletion (compare Fig. 1d with Fig. 5d). Upregulation of Xist RNA upon acute depletion of ZCCHC8/NEXT is evident across the entire transcript (Extended Data Fig. 8c), and principal component analysis indicates that accelerated silencing affects X-linked genes equivalently across the X chromosome (Extended Data Fig. 8d). Levels of Kcnq1ot1 RNA were similarly elevated following acute depletion of ZCCHC8 but not ZFC3H1 (Extended Data Fig. 3g-i, right). SLAM-seq analysis indicates that elevated Xist RNA levels following acute ZCCHC8 depletion are attributable to increased transcript stability (Extended Data Fig. 4c). Taken together these results suggest that m^6^A promotes Xist RNA turnover by the NEXT complex independently of YTHDC1.

## Discussion

The acceleration of Xist-mediated silencing that we observe following acute METTL3 depletion conflicts with prior studies that used long-term knockout/knockdown strategies and reported abrogation of silencing to a greater or lesser degree^16,17^. Differences observed in our experiments are likely attributable to reduced indirect effects from acute as opposed to chronic depletion of METTL3. This conclusion is supported by the observation that global mRNA expression changes show an improved correlation with m^6^A target mRNAs with acute METTL3 depletion compared to chronic knockout/knockdown. Although we cannot pinpoint with certainty why perturbation of the m^6^A pathway in prior studies resulted in abrogated Xist-mediated silencing, a possible explanation could lie in changed levels of mRNAs encoding proteins which regulate expression of key silencing factors. Indeed, mRNAs encoding the silencing factors SPEN, HDAC3, and HNRNPU are all m^6^A modified. Regardless, our findings underscore that caution needs to be exercised in interpreting experiments involving chronic perturbation of the m^6^A pathway due to its roles in regulating transcription/RNA metabolism at a global level^34,35^.

Our results indicate that accelerated silencing following acute METTL3 depletion is linked to increased levels of Xist RNA which in turn result from decreased Xist RNA turnover. This interpretation is supported by the similar phenotype seen following acute depletion of the NEXT subunit ZCCHC8. We envisage that higher Xist RNA levels result in an increased number of molecules accumulating on the X chromosome, thereby amplifying the recruitment of silencing factors that mediate gene silencing in X inactivation. Of note, the increased Xist RNA levels linked to reduced turnover are tempered by a reduction of Xist transcription rates, consistent with our prior analyses indicating feedback control between Xist transcription and stability^25^. Accordingly, we predict that perturbation of the feedback control pathway(s) that inhibit Xist transcription should similarly enhance Xist RNA levels and accelerate the rate of silencing.

That NEXT but not PAXT regulates Xist RNA turnover is perhaps unexpected given that Xist is polyadenylated and NEXT has been linked to regulating non-polyadenylated nuclear RNAs such as PROMPTs and eRNAs^36^. One possible explanation is that the poly(A) tail of Xist molecules localised along the Xi chromosome is protected from NEXT activity, for example by poly(A) binding proteins such as PABPN1 or by a folded back structure similar to the lncRNA MALAT1^37^. Progressive erosion of this protection over time would thus initiate NEXT-exosome complex engagement to trigger degradation from the 3’ end. Consistent with this idea our previous RNA-SPLIT analysis demonstrates that Xist molecules in WT cells remain relatively stable for an extended period of around 90 minutes before showing rapid exponential degradation kinetics^25^.

The marked increase in Xist RNA stability, Xist RNA levels and Xi silencing rate observed with both METTL3 and ZCCHC8 depletion, suggests that m^6^A and the NEXT complex function together to regulate Xist RNA turnover. Based on prior studies identifying the YTHDC1-NEXT axis and its role in regulating carRNAs, we anticipated that YTHDC1 could bridge Xist RNA with NEXT for m^6^A-mediated control of turnover. However, given that acute YTHDC1 depletion has no significant effect on Xist RNA turnover, we speculate that there is an alternative pathway that links m^6^A to the NEXT complex. In support of this idea, we observe identical YTHDC1 independent effects on levels of Kcnq1ot1, which like Xist is chromatin associated and functions to silence genes in *cis*^38^. We envisage two possible models, a direct pathway whereby an unidentified reader protein interacts both with m^6^A and NEXT, or an indirect pathway where m^6^A promotes association of Xist RNA-binding proteins that inhibit NEXT/exosome activity (Fig. 5g). In relation to the latter possibility, hnRNPC and hnRNPA2B1 proteins that have been previously reported to function indirectly via an “m^6^A-switch” mechanism^39,40^ were identified in mass spectrometry experiments as Xist RNA interactors^8^.

In conclusion, we find that the m^6^A pathway functions in conjunction with the NEXT complex to modulate Xist RNA turnover, and thereby contributes to the feedback control mechanisms that determine Xist RNA levels during normal development.

## Acknowledgement

We are grateful to Flavia Constantinescu for assistance with mESC tissue culture during pandemic, to Joseph Bowness for helping with the initial RNA-seq library preparation, to Adam Cawte for helping with construct design for knock-in at Rosa26 locus, to Yixin Zhang for quantifying site-specific m^6^A changes on Xist, and to all the Brockdorff lab members for fruitful discussions. We thank Oxford Biochemistry IT support for computing server maintenance. We thank Benoît Moindrot for critical reading of the manuscript. Work in the Brockdorff lab is supported by Wellcome Trust (215513/Z/19/Z) and UKRI (EP/Y029062/1).

## Author contributions

G.W. and N.B. conceived the study. G.W., with assistance from M.A., L.R., and T.B.N., generated the dTAG degron cell lines. G.W. conducted the ChrRNA-seq, SLAM-seq, total RNA-seq, and performed all sequencing data analyses. L.R., under the supervision of H.C. and N.B., developed RNA-SPLIT and applied it to Xist in this study. H.C., H.L.R., and L.R. analyzed the RNA-SPLIT data and produced the plots. N.B. and G.W. drafted the manuscript with contributions from co-authors. All authors read and approved the manuscript. N.B. supervised the entire project.

## Competing interests

The authors declare no competing interests.

## Methods

### Tissue Culture

All mouse embryonic stem cells (mESCs) were grown in feeder-dependent conditions on gelatinized plates at 37°C in a 5% CO_2_ incubator. Mitomycin C-inactivated mouse fibroblasts were used as feeders. mESCs medium consists of Dulbecco’s Modified Eagle Medium (DMEM, ThermoFisher) supplemented with 10% foetal calf serum (Merck), 2 mM L-glutamine (ThermoFisher), 1× non-essential amino acids (ThermoFisher), 50 μM *β*-mercaptoethanol (ThermoFisher), 50 g/mL penicillin/streptomycin (ThermoFisher), and 1 mL of Leukemia Inhibitory Factor (LIF)-conditioned medium made in-house. *Xist* expression was induced by the addition of 1 µg/mL doxycycline (Dox) (Sigma-Aldrich, D9891) for 24 hours. Protein of interest with FKBP12^F36V^ knock-in was trigged for degradation by the addition of dTAG-13 (100 nM) (gift from James Bradner lab).

### Chromatin-associated RNA-seq (ChrRNA-seq)

Chromatin associated RNA was extracted according to Nesterova et al. 2019 (ref^17^). Briefly, mESCs from one confluent 15 cm dish were trypsinised and washed in PBS. Cells were lysed on cold ice in RLB buffer [10 mM Tris pH 7.5, 10 mM KCl, 1.5 mM MgCl_2_, and 0.1% NP40], and nuclei were purified by centrifugation through a sucrose cushion (24% sucrose in RLB). Pelleted nuclei were resuspended in NUN1 [20 mM Tris-HCl pH 7.5, 75 mM NaCl, 0.5 mM EDTA, 50% glycerol], then lysed with NUN2 [20 mM HEPES pH 7.9, 300 mM, 7.5 mM MgCl_2_, 0.2 mM EDTA, 1 M Urea]. Samples were incubated for 15 minutes on ice with occasional shaking, then centrifuged at 2800 g to isolate the insoluble chromatin fraction. The chromatin pellet was resuspended in TRIzol by passing through a 23 G needle several times. Finally, chromatin-associated RNA was purified through standard TRIzol/chloroform extraction followed by isopropanol precipitation. Samples were treated with 2 rounds of Turbo DNaseI to remove the DNA contamination. The quality of the ChrRNAs were checked by RNA bioanalyzer. Approximately 250-750 ng of RNA was used for library preparation using the Illumina TruSeq stranded total RNA kit including the rRNA depletion step (RS-122-2301). Libraries were quantified by qPCR with KAPA Library Quantification DNA standards (Kapa biosystems, KK4903). DNA and RNA concentrations were also determined by Qubit. The libraries were pooled and 2x 81 paired end sequencing was performed using Illumina NextSeq 500 (FC-404-2002).

### Western Blot

Total cell lysates or quantified cell fractionations were resolved on a polyacrylamide gel and transferred onto PVDF or nitrocellulose membrane by quick transfer. Membranes were blocked by incubating them for 1 hour at room temperature in 10 mL 5% w/v Marvell milk powder. Blots were incubated overnight at 4°C with the primary antibody, washed 4 times for 10 minutes with PBST and incubated for 1 hour with secondary antibody conjugated to horseradish peroxidase. After washing 5 times for 5 min with PBST, bands were visualised using ECL (GE Healthcare). TBP, SETDB1, KAP1 serve as loading controls. Antibodies used in this study are: METTL3 (Abcam, ab195352), METTL14 (Sigma-Aldrich, HPA038002), RBM15 (Proteintech, 10587-1-AP), YTHDC1 (Sigma-Aldrich, HPA036462), TBP (Abcam, ab51841), SETDB1 (Proteintech, 11231-1-AP), ZCCHC8 (Proteintech, 23374-1-AP), ZFC3H1 (Sigma-Aldrich, HPA007151), and KAP1 (Abcam, ab10484).

### Molecular cloning and CRISPR-Cas9 mediated knock-in

We followed an established strategy for FKBP12^F36V^ knock-in^21^. Briefly, sgRNA targeting near *Ythdc1 or Zcchc8* stop-codon was designed by online tool CRISPOR^41^ (http://crispor.tefor.net/) and oligos were synthesised from Invitrogen, then cloned into pSpCas9(BB)-2A-Puro (PX459) V2.0 (Addgene #62988) backbone following instructions. A donor vector was built by Gibson assembly (NEB) of homology arms (∼400 bp) PCR amplified from genomic DNA and the FKBP12^F36V^ sequence amplified from plasmid pLEX_305-N-dTAG (Addgene #91797)^20^, or other coding sequences (e.g. GFP-METTL3) described below. To achieve site-specific mutagenesis from the plasmid, we followed the protocol from QuikChange Lightning Site-Directed Mutagenesis Kit (Agilent) or Gibson Assembly. This strategy was used for generating the mESC line with GFP-METTL3 or GFP-METTL3 with D395A at *Rosa26* locus (see below). The Cas9-sgRNA-containing plasmid and donor vector were co-transfected at a molar ratio of 1:6 into XX mESCs cells on a 6-well plate using Lipofectamine 2000 according to the manufacturer’s protocol (ThermoFisher).

Transfected cells were passaged at different densities into three Petri dishes with feeders, 24 h after transfection. The next day, cells were subjected to puromycin selection (4.5 μg/mL) for 48 h and then grown in regular mESC medium until mESC colonies were ready to be picked and expanded. Western blot analysis was used to screen colonies for FKBP12^F36V^ knock-in as this results in slower mobility of the FKBP12^F36V^ fused protein compared to the wild-type protein in an SDS-PAGE gel and disappearance of the WT band on western blot. Selected clones were further characterized by PCR of genomic DNA and Sanger sequencing to confirm correct knock-in and homozygosity. The karyotype status of the X chromosome was checked by PCR. Primers used in this study are listed in Supplementary Table X. The sensitivity of selected clones to dTAG-13 treatment was validated by western blot.

### Complementation using METTL3 or catalytic mutant METTL3 transgenes in the *Rosa26* locus

CRISPR-Cas9 mediated knock-in was used to integrate METTL3 or catalytic mutant METTL3 (D395A)^24^ transgenes into the *Rosa26* locus. In order to distinguish transgene encoded from endogenous METTL3 and FKBP12^F36V^ tagged METTL3, we added sequence encoding an in frame N-terminal GFP tag. sgRNA (<u>CGCCCATCTTCTAGAAAGAC</u>) was used for targetted knock-in. D395A mutated METTL3 was constructed by Gibson Assembly. PCR and Western blot were used for screening colonies. The expression and nuclear localisation of the GFP-METTL3 fusion proteins were confirmed by fluorescence microscopy. Elimination of the X_129_ chromosome occurred in several clones as (as indicated in figurelegends). This did not preclude assaying relative silencing efficiency and Xist RNA levels in the presence and absence of doxycycline and dTAG-13.

### RNA-SPLIT

RNA-SPLIT was carried out as detailed previously^25^. Briefly, mESCs were grown on gelatin-coated 18×18 mm No 1.5H precision coverslips (± 5 μm tol.; Marienfeld Superior) in a 6-well plate on a layer of feeder cells. When the mESCs reached 60-70% confluency, Xist expression was induced for 1.5 or 24 h using 1 μg/mL doxycycline. Cells were then incubated with 50 nM diAcFAM HaloTag ligand (488 nm, Promega) and 1 μg/mL doxycycline for 45 min, before being washed with ESC medium containing 1 μg/mL doxycycline for 15 min. Next, different coverslips were incubated with 50 nM JF-585 HaloTag ligand (kindly provided by Luke Lavis, HHMI Janelia) and 1 μg/mL doxycycline for different times to label newly synthesized Xist RNA molecules, (0, 60, 80, 100, 120, 140, 160, 180, 200, and 220 min to assess Xist turnover or 10, 20, 30, 40, 50, and 60 min to assess Xist RNA transcription dynamics) before being washed with PBS. Cells were fixed for 10 min at room temperature with 2% formaldehyde prepared fresh in PBS (pH 7) before a stepwise exchange to PBST (0.05% Tween) and permeabilization with 0.2% Triton X-100 for 10 min, followed by two washes with PBST. Subsequently, cells were incubated with 2 μg/mL DAPI in PBST for 10 min, before being washed briefly with PBS, followed by ddH2O, and mounted centrally on the unfrosted side of Superfrost Plus microscopy slides (VWR) using Vectashield soft mount media, sealed with clear nail polish and imaged using the DeltaVision OMX V3 Blaze system (GE Healthcare). Images were analysed as described previously^25^, with all scripts and details of further script refinement to improve usability as detailed.

### Total RNA-seq (RNA-seq)

METTL3-FKBP12^F36V^ taggedcells (C3 and H5) were either treated with dTAG-13 for 26 hours or not. Cells were washed with PBS twice and directly lysed with Trizol, followed by total RNAs isolation as the aforementioned ChrRNA-seq. DNA contamination was removed by Turbo DNase I treatment. The quality of the RNAs were checked by RNA bioanalyzer. Approximately 500 ng of total RNA was used for library preparation using the Illumina TruSeq stranded total RNA kit including the rRNA depletion step (RS-122-2301). The libraries were pooled and 2x 81 paired end sequencing was performed using Illumina NextSeq 500 (FC-404-2002).

### SLAM-seq

SLAM-seq was performed using the SLAMseq Kinetics Kit Catabolic Kinetics Module (cat. no. 062.24, Lexogen) as previously described^25^ and also shown as schematic in Extended Data Fig. 4a. Specifically, cells were grown in gelatin-coated 6-well plates after pre-plating to discard feeder cells. When reaching 60 to 70% confluency, cells were either untreated or dTAG-13 treated for 2 hours, followed by additional 1 μg/ml doxycycline induction for 20 hours. Transcribed RNA was then labelled with 4sU by incubation with medium containing 500 μM 4sU (Lexogen) and 1 μg/ml doxycycline for another 4 hours. 4sU was withdrawn for all samples by medium washout. Different samples were washed with medium containing 1 μg/ml doxycycline and 50 mM uridine (in excess, Lexogen) for 1.5 and 3 hours. Cells were washed with PBS once and directly lysed with Trizol/chloroform for total RNA isolation. Equal amounts of RNA (5 μg) were treated with iodoacetamide to modify the 4-thiol group of S4U-containing nucleotides via the addition of a carboxyamidomethyl group by the SLAMseq Kinetics Kit (Lexogen). The RNA was recovered using RNAClean XP beads (Beckman Coulter), followed by resuspension in nuclease free water. Approximately 500 ng RNA of each sample was taken forward for library preparation using the Illumina TruSeq stranded total RNA kit (RS-122-2301). Quantification of the libraries was conducted by qPCR using KAPA Library Quantification DNA standards (Kapa Biosystems, KK4903). Finally, the libraries were pooled and 2× 81 paired-end sequencing was performed using Illumina NextSeq500 (FC-404-2002).

### SLAM-seq data analysis

Estimation of RNA half-life was performed using GRAND-SLAM^42^. Briefly, the paired-end sequencing reads were first aligned to rRNA, and then the unmapped reads were mapped to mouse genome mm10 by STAR (v2.5.2b)^43^ with the key parameters (*--twopassMode Basic -- outSAMstrandField intronMotif --outSAMattributes All --outFilterMultimapNmax 1 -- outFilterMismatchNoverReadLmax 0.06 --alignEndsType EndToEnd*). Given that T to C conversion is the signature of SLAMseq, the T to C conversion rate was calculated as 4sU incorporation and the average of all the rest of the conversions was calculated as background due to errors from sequencing or library preparation. The T to C conversion rate and background rate were calculated accordingly for each sample. The RNA decay was assumed to follow an exponential model, so the corresponding background corrected T2C rates were fitted to the exponential model to estimate Xist RNA half-life.

### ChrRNA-seq data analysis

The ChrRNA-seq data mapping pipeline used in this study was similar to previous work^17^. Briefly, the raw fastq files of read pairs were first mapped to an rRNA build by bowtie2 (v2.3.5.1)^44^ and rRNA-mapped reads discarded. The remaining unmapped reads were aligned to the ‘N-masked’ genome (mm10) with STAR (v2.5.2b)^43^ using parameters (*--twopassMode Basic -- outSAMstrandField intronMotif --outFilterMismatchNoverReadLmax 0.06 --outFilterMultimapNmax 1 --alignEndsType EndToEnd*) for all the sequencing libraries. Unique alignments were retained for further analysis. We made use of 23,005,850 SNPs between Cast and 129S genomes and employed SNPsplit (v0.4.0dev, Babraham Institute, Cambridge, UK) to split the alignment into distinct alleles (Cast and 129S) using the parameter “*--paired*”. The (allelic) read numbers were counted by the program featureCounts (*-t transcript -g gene_id -s* 2)^45^ and the alignments were sorted by Samtools^46^. Files of bigWig were generated by Bedtools^47^ and visualized by IGV^48^ or UCSC Genome Browser. Metagene profile and heatmap were generated by deepTools^49^, as well as custom python scripts. For biallelic analysis, counts were normalized to 1 million mapped read pairs (as CPM) by the R package (v4.1.0) of *edgeR*. Genes with at least 10 SNP-covering reads across all the samples were used to calculate the allelic ratio of Xi/(Xi+Xa) where Xi and Xa indicate inactive and active allele, respectively. Principal components analysis (PCA) was done using the *prcomp* function in R and plotted using the ggplot2 package. Allelic ratio of samples derived from SPEN KO or deletion of Xist B/C-repeat were taken from the previous study^17^. Gene categories including initial X-linked gene expression level and promoter chromatin landscape were taken from Nesterova et al, 2019 and Pintacuda et al 2017, gene silencing kinetics data were taken from Bowness et al, 2022.

### Total RNA-seq data analysis

Total RNA-seq data analysis procedure was similar to ChrRNA-seq regarding the raw fastq reads mapping. PCR duplicates were removed by Picard command line, MarkDuplicates. Differentially expressed genes and transposable elements were called by TEtranscripts^50^ (v2.2.1). Up and down regulated genes called were further cross compared with m^6^A annotations from previous studies^21,51^.

## Data availability

High-throughput raw sequencing data as well as key processed data, including ChrRNA-seq, SLAM-seq, and total RNA-seq, are deposited to the National Center for Biotechnology Information’s Gene Expression Omnibus (accession number GSEXXXXX). Imaging data are available from XXXXX.

## Code availability

Key parameters used for sequencing data analysis are detailed in the above methods. The RNA-SPLIT analysis code was deposited to GitHub (https://github.com/HollyRoach/Automated_RNA-SPLIT).

## Extended Data Figure

**Extended Data Fig. 1:**
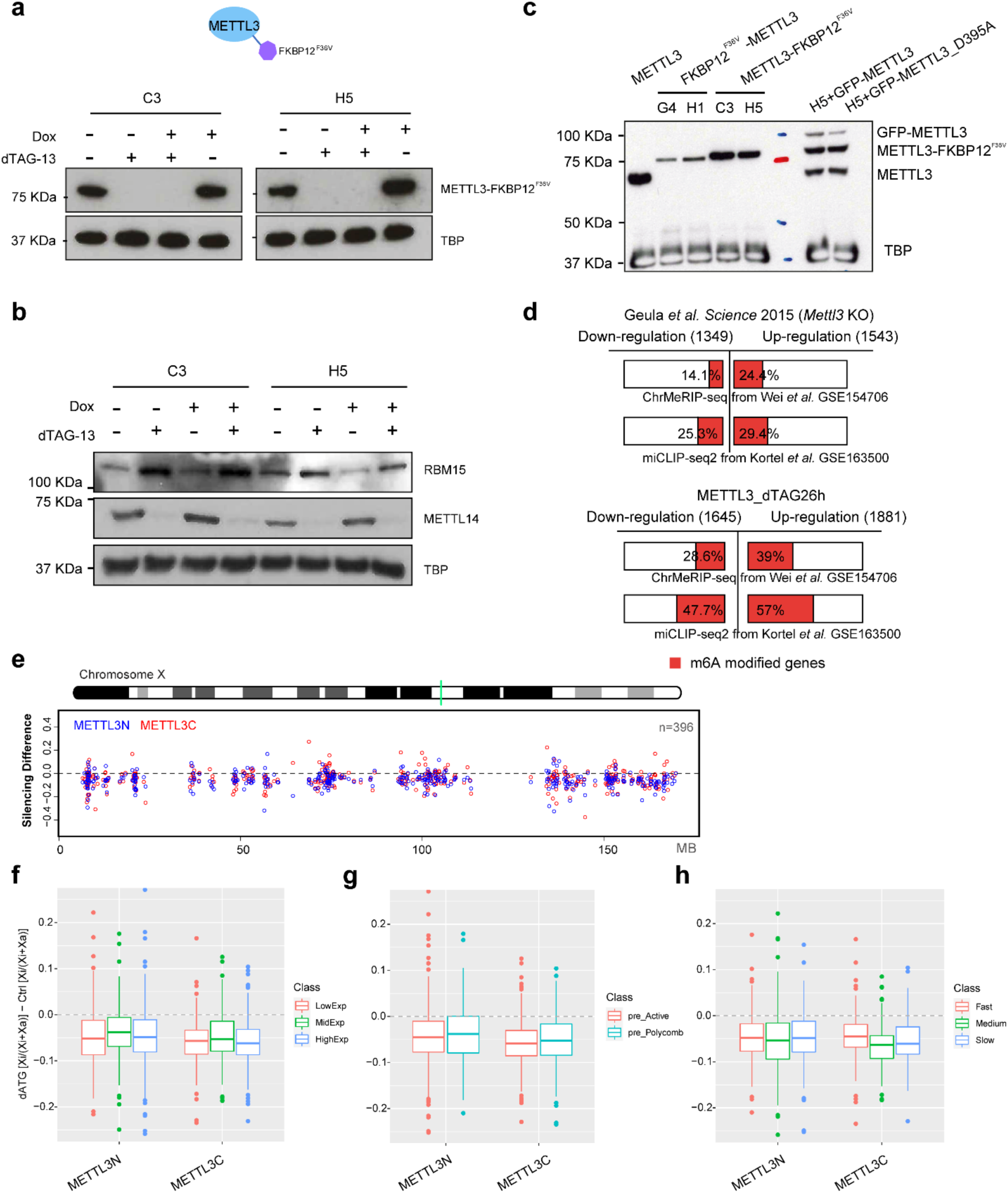
Derivation of degron tagged METTL3 cells and their effect on Xist-mediated silencing. **a**, Western blot showing the METTL3-FKBP12^F36V^ protein level in two independent clones (C3 and H5) under ChrRNA-seq analysis conditions shown in Fig. 1d . TBP loading control. **b**, Western blot showing the protein level for RBM15 and METTL14 for C3 and H5 clones under ChrRNA-seq analysis conditions shown in Fig. 1d. **c**, Western blot showing protein size and level for WT METTL3, FKBP12^F36V^ tagged METTL3, and GFP-tagged WT or catalytic mutant METTL3. TBP loading control. **d**, Bar plot showing the proportion of differentially expressed genes in *Mettl3* KO mESCs^23^ (top) or 26 hours dTAG-13 treated METTL3-FKBP12^F36V^ mESCs (bottom) with m^6^A modification. The number of differentially up- or down-regulated genes are indicated. m^6^A peaks/sites are annotated from ChrMeRIP-seq data^21^ and miCLIP-seq2 data^51^. **e**, Allelic ratio difference between dTAG + Dox samples and Dox samples in Fig. 1d. Values from METTL3 N-terminal or C-terminal FKBP12^F36V^ are averaged and shown as METTL3N (blue) and METTL3C (red) respectively. Green line on ChrX ideogram indicates location of Xist. Dashed line denotes 0, indicating no silencing difference. **f, g, h**, As in (e), but for gene group analysis. Boxplots showing the difference of allelic ratio towards three different gene categories defined previously;initial X-linked gene expression level^17^ (f), initial promoter chromatin state^15^ (g), gene silencing kinetics^22^ (h). FKBP12^F36V^ indicated as dTAG.

**Extended Data Fig. 2:**
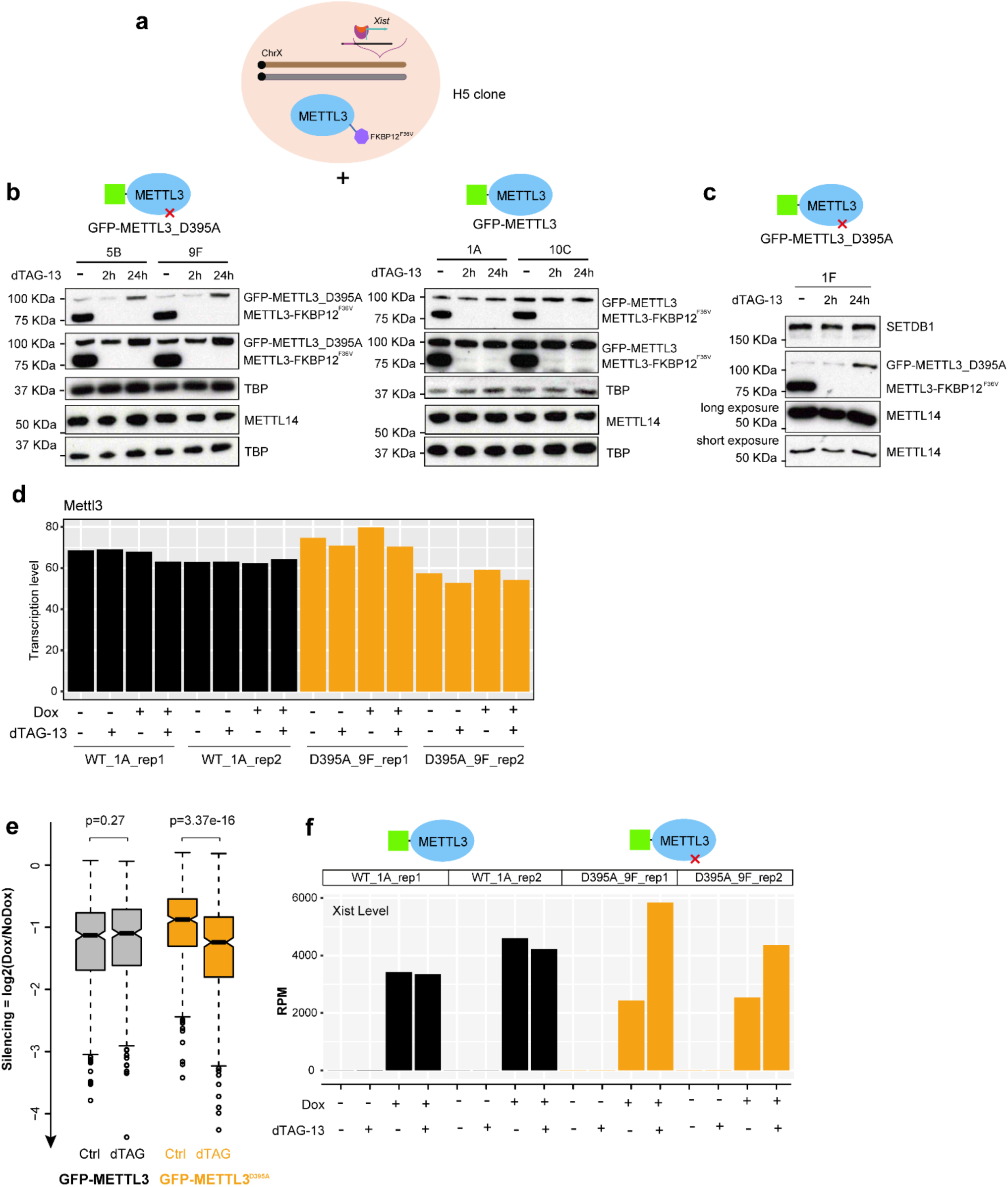
METTL3 catalytic activity is required for acceleration of Xist-mediated silencing. **a**, Schematic showing the strategy for transgene complementation. METTL3 transgene is inserted into *Rosa26* locus and transcribed under the native *Rosa26* promoter. **b,c** Western blots showing the protein level for GFP-METTL3^D395A^ (left: 5B, 9F, 1F are three independent clones) and GFP-METTL3 (right: 1A and 10C are two independent clones), along with METTL3-FKBP12^F36V^ and METTL14, after 2 or 24 hours dTAG-13 treatment. TBP and SETDB1 loading controls. Note that clones 5B, 9F, 1A, and 10C are X_cast_0 due to loss of X_129_. Clone 1F has retained both X_129_ and X_cast_. Here X_cast_ is the Xi chromosome. **d,** Barplot showing Mettl3 transcript level based on ChrRNA-seq data for cells harbouring GFP-METTL3 (represented by clone 1A) or GFP-METTL3D^395A^ (represented by clone 9F). Two replicates for each clone are included. Expression level is the sum of reads from native *Mettl3* locus and transgenic *Rosa26* locus. The yellow bars represent the level from cells expressing the transgene encoding METTL3^D395A^. **e**, Boxplot showing average gene expression levels for X-linked genes in control or dTAG-13 treated samples. The averages are calculated from two clones shown in (b). Yellow boxplots represent the level from cells expressing the transgene encoding METTL3^D395A^. **f**, Xist expression level from samples of GFP-METTL3 or GFP-METTL3^D395A^ clones. Clone names are indicated as above. Y-axis denotes the RPM value, as shown in Fig. 2a. Yellow boxplots represent the level from cells expressing the transgene encoding METTL3^D395A^.

**Extended Data Fig. 3:**
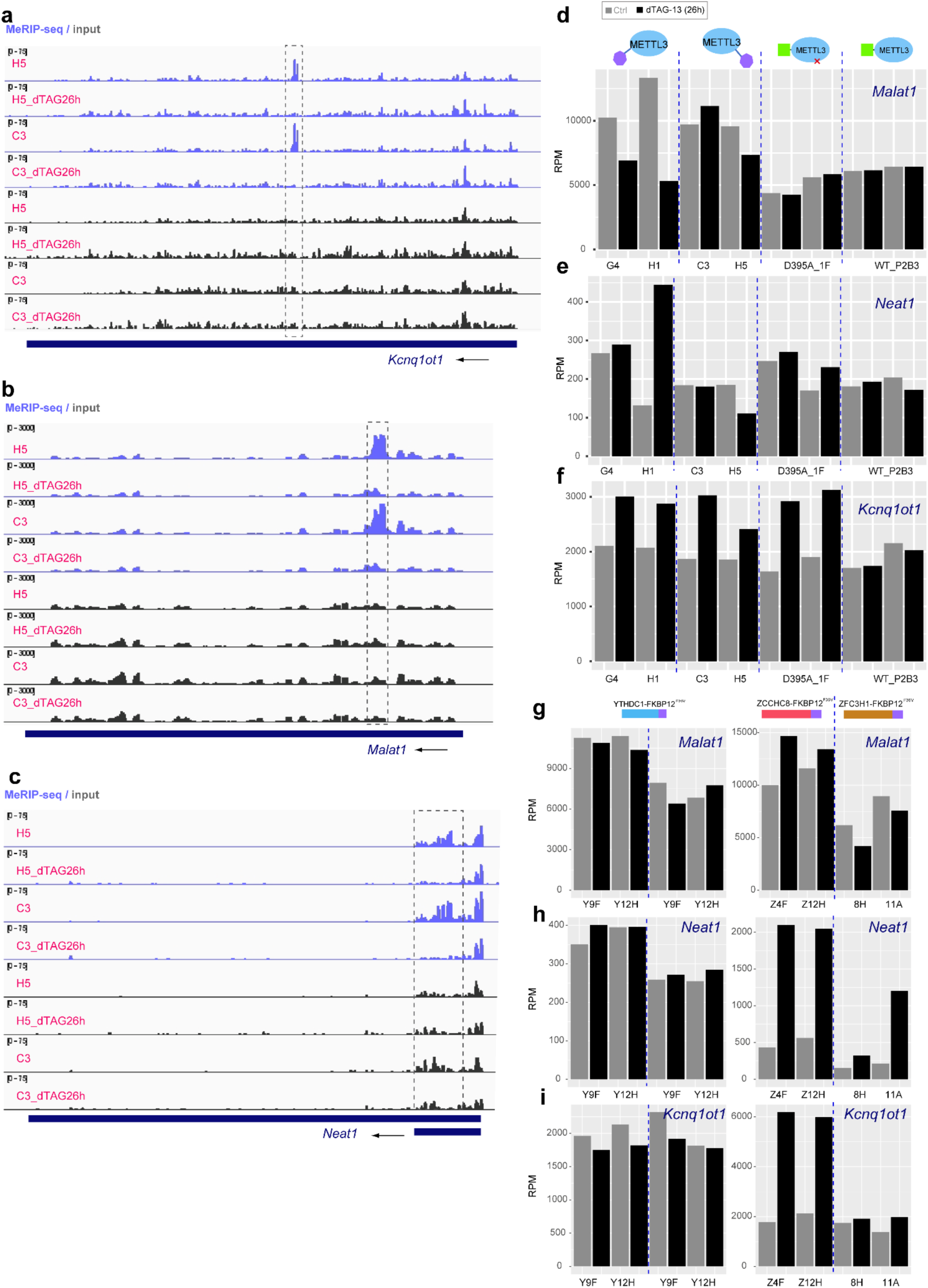
m^6^A landscape and gene expression analysis for lncRNAs Malat1, Neat1, and Kcnq1ot1. **a-c**, m^6^A profile on lncRNAs Kcnq1ot1 (**a**), Malat1 (**b**), and Neat1 (**c**). Data are from ref^21^. MeRIP-seq data is shown in blue and input data in grey. For each panel, gene structure and transcriptional direction are shown below. Black boxes highlight METTL3 dependent m^6^A modification peaks. **d-f**, Expression levels for lncRNAs Malat1 (**d**), Neat1 (**e**), and Kcnq1ot1 (**f**) from ChrRNA-seq analysis. Cell lines are indicated at the bottom. Samples are from METTL3-FKBP12^F36V^ and transgene complementation experiments. Control samples are shown in grey, whereas the dTAG-13 treated (26 hours) samples are shown in black. **h-i**, As in (**d-f**), but for samples YTHDC1-FKBP12^F36V^ (left), ZCCHC8-FKBP12^F36V^ (right), and ZFC3H1-FKBP12^F36V^ (right). Note that two replicates are shown for each YTHDC1 clone.

**Extended Data Fig. 4:**
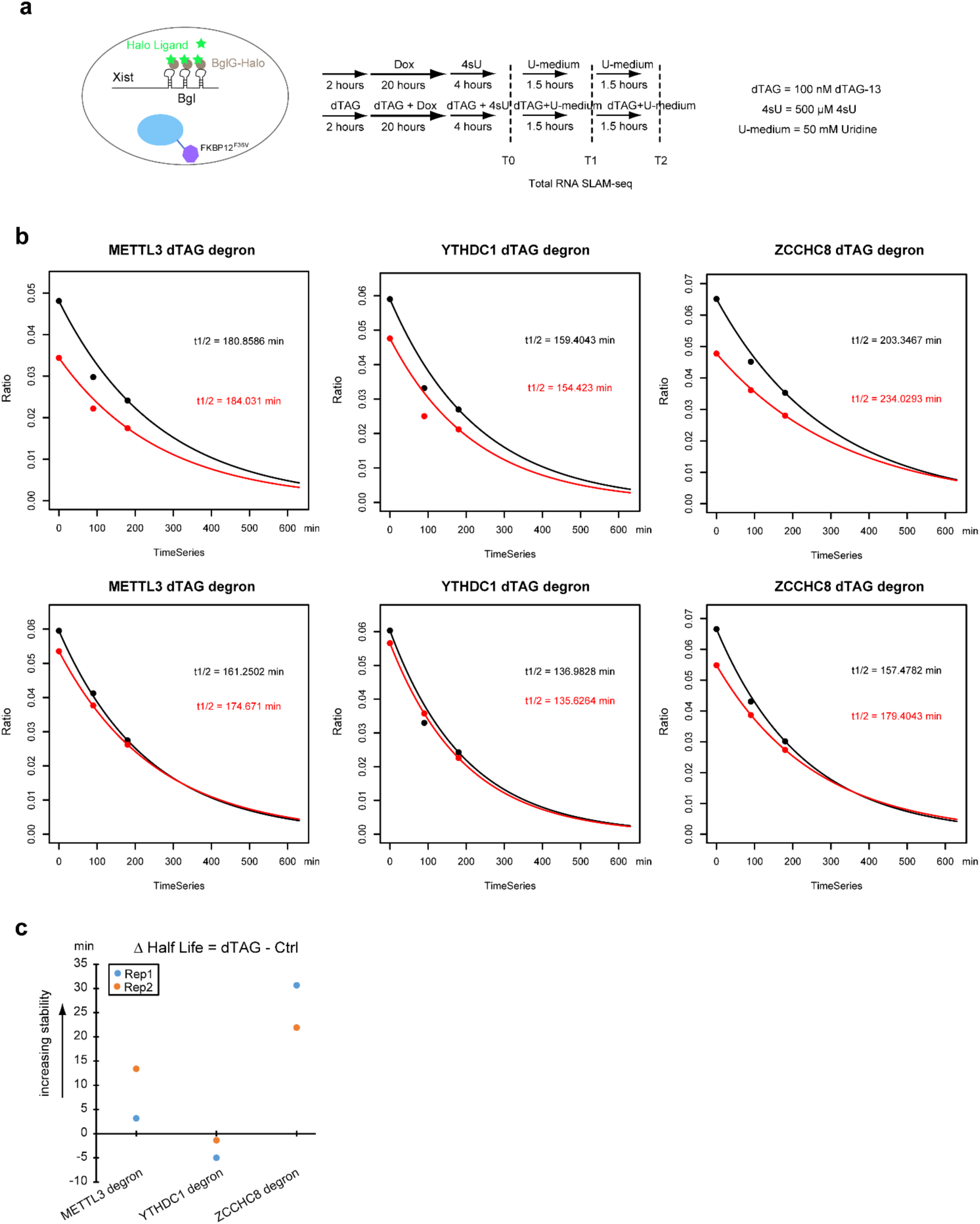
Xist half-life estimated by SLAM-seq analysis. **a,** Schematic showing the experimental design for SLAM-seq. Cells used in this analysis are in the Xist-BglG expressing lines with -FKBP12^F36V^ tags on METTL3, YTHDC1, or ZCCHC8. **b**, Best fit curves indicating Xist degradation determined from SLAM-seq T-to-C conversions. Two independent experiments are shown (top and bottom panels). Black and red represent untreated and dTAG-13 treated samples respectively. **c**, Dot plot showing the difference of Xist half-life with and without dTAG-13 treatment from (b). Replicates 1 and 2 are shown in blue and orange respectively.

**Extended Data Fig. 5:**
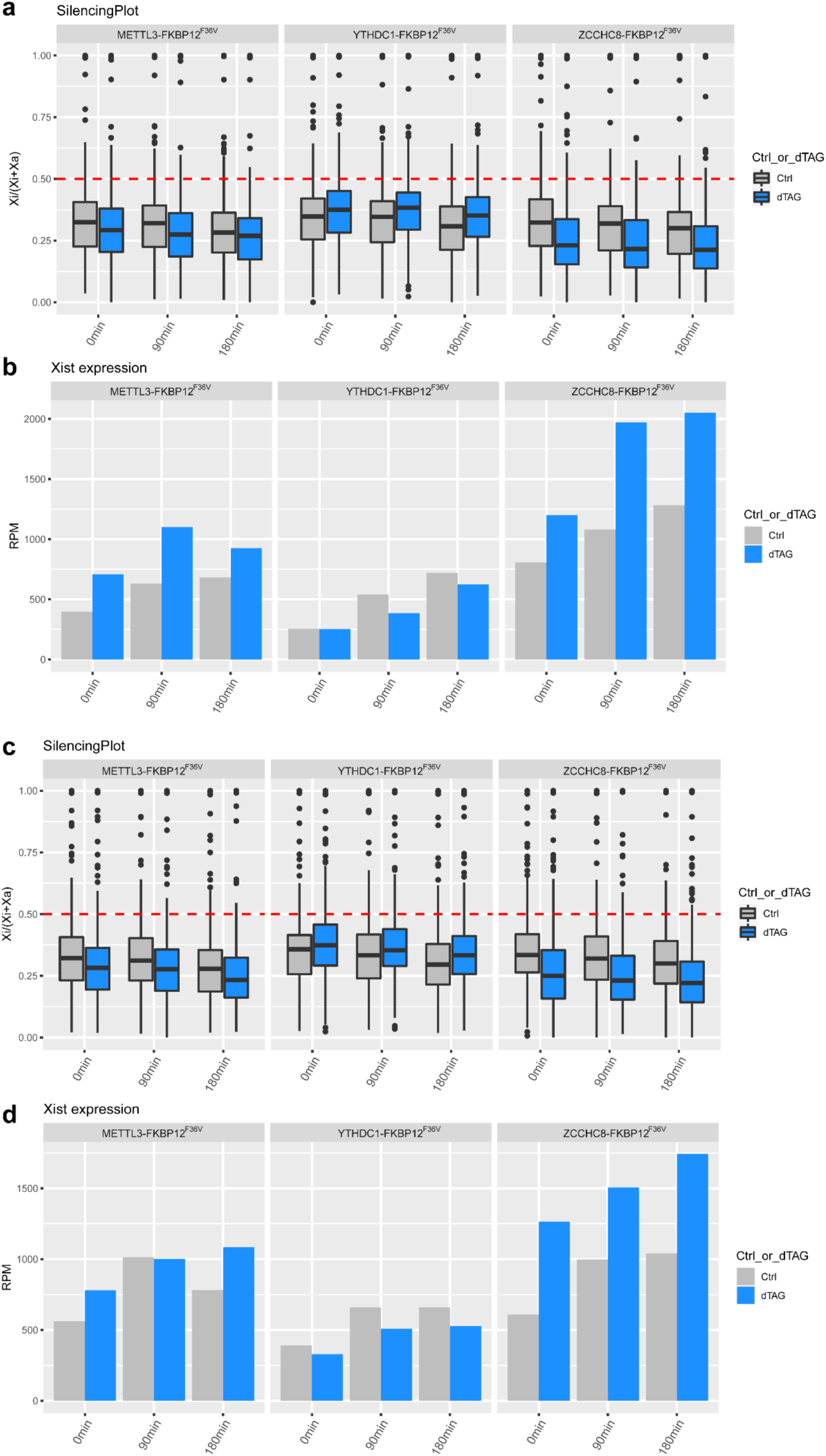
X-inactivation dynamics and Xist levels calculated from SLAM-seq experiments. **a**, Boxplot showing X-inactivation dynamics in the SLAM-seq timecourse experiment shown in Extended Data Fig. 4. Grey and blue indicate untreated and dTAG-13 treated samples respectively. The dotted line indicates allelic ratio of 0.5. **b**, Xist expression level in the samples in (a). **c,d**, As in a,b for replicate 2 experiment.

**Extended Data Fig. 6:**
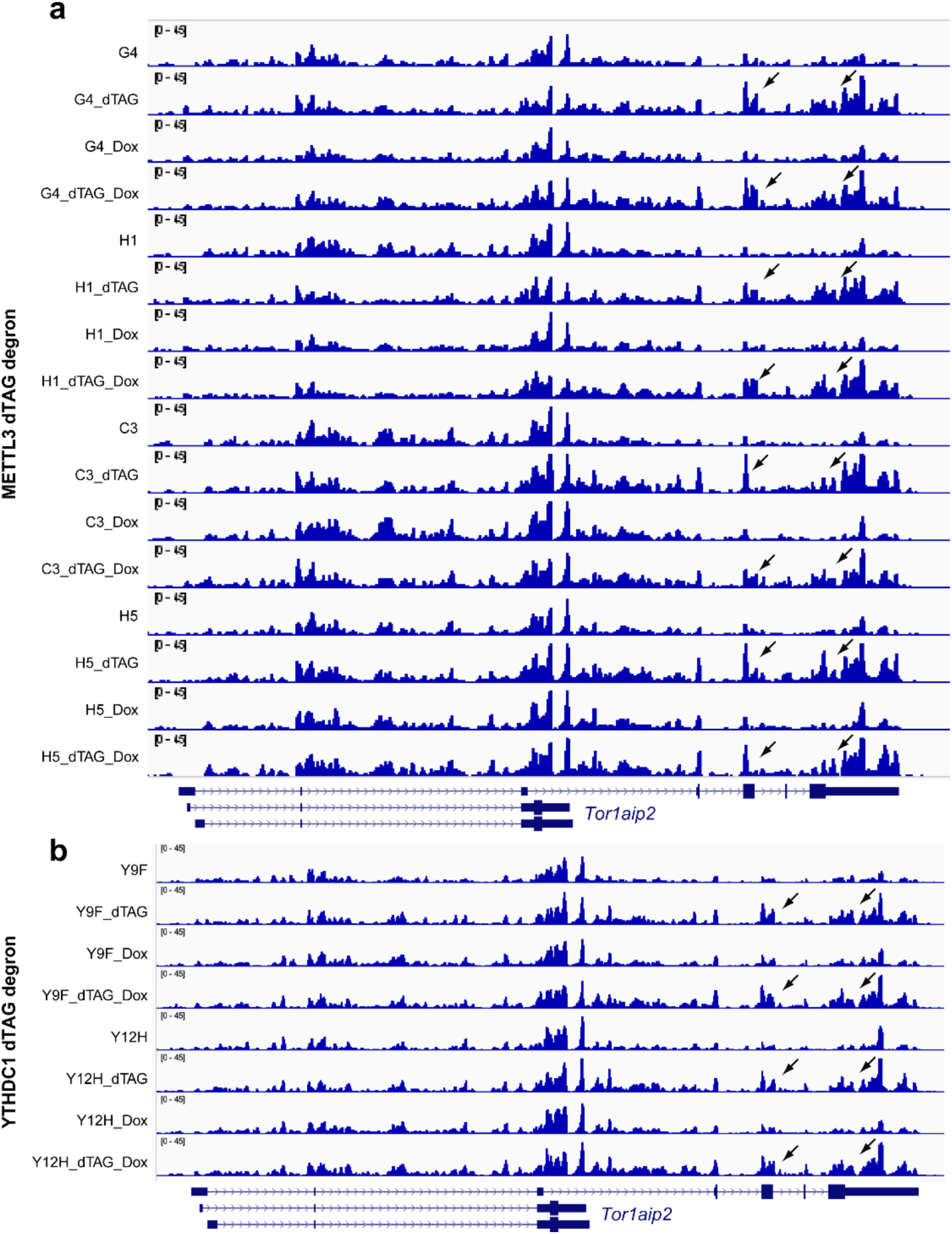
FKBP12^F36V^ degron for nuclear m^6^A reader protein YTHDC1. **a,b**, Profile of Tor1aip2 short and long isoforms in ChrRNA-seq samples of (a) METTL3-FKBP12^F36V^ cells and (b) YTHDC1-FKBP12^F36V^ degron cells (b). The arrows indicate the long isoform strongly upregulated in all of the dTAG-13 treated samples compared to untreated samples. Tor1aip2 gene structure is shown below.

**Extended Data Fig. 7:**
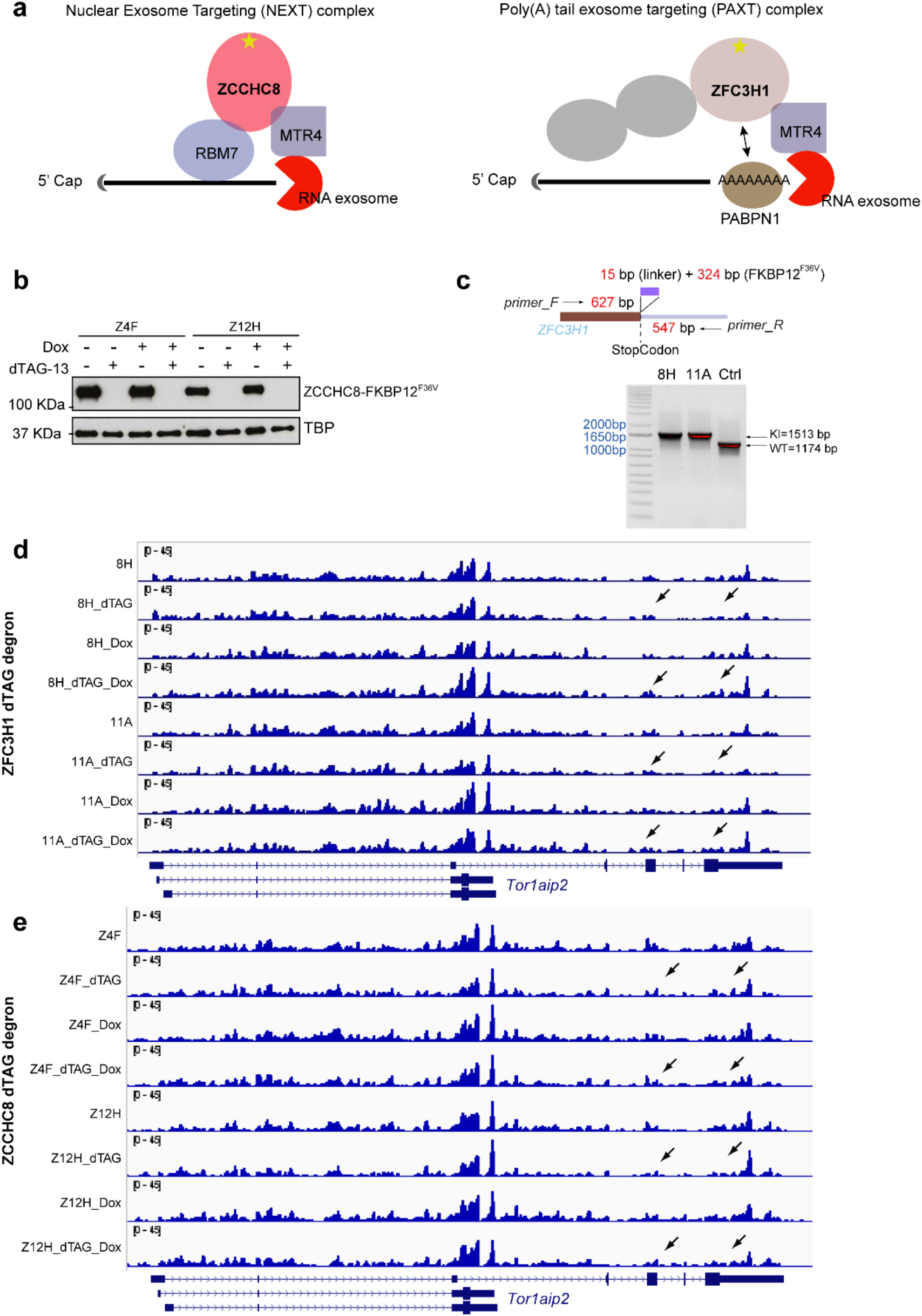
dTAG degron for ZFC3H1, a core component of PAXT complex, and ZCCHC8, a core component of NEXT complex. **a,** Schematics illustrating the action of NEXT and PAXTcomplexes for nuclear RNA degradation. FKBP12^F36V^ tagged subunit is indicated (star). **b**, Western blot verifying selected clones Z4F and Z12H in Fig. 5c. TBP loading control. **c**, Validation of FKBP12^F36V^ insertion in the selected clones in Fig. 5b,d using PCR. **d,e**, Profile of Tor1aip2 short and long isoforms in ChrRNA-seq samples shown in Fig. 5c,d. The arrows indicate that the long isoform is not upregulated in dTAG-13 treated samples compared to untreated samples. See also Extended Data Fig. 6. Tor1aip2 gene structure is shown below.

**Extended Data Fig. 8:**
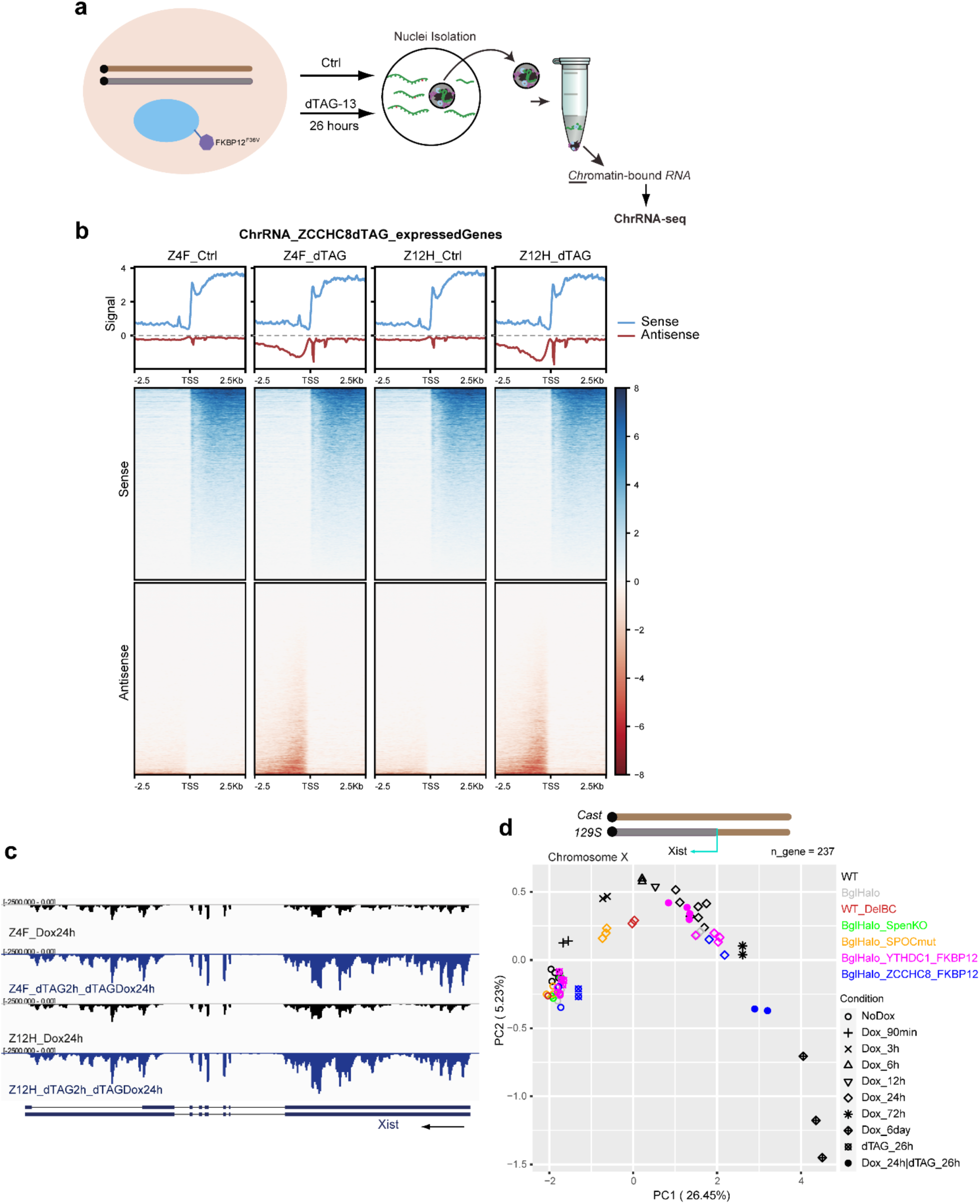
Expression profile of PROMPTs, Xist RNA, and X-linked silencing in ZCCHC8-FKBP12^F36V^ cells. **a**, Schematic showing the experimental procedure for control or 26 hours dTAG-13 treated samples for ChrRNA-seq analysis. **b**, Aggregated profile (top) and heatmap (bottom) showing the expression of expressed genes (blue) and their promoter upstream transcripts (PROMPTs) (red). The negative value shown in red in this plot indicates antisense transcription. Plots show data for transcription start site ± 2.5 kb. **c**, Genome browser screenshot of ChrRNA-seq of the Xist gene from ZCCHC8 degron samples with Dox or dTAG-13 + Dox treatment. **d**, PCA using allelic ratio of X-linked genes (n=237) for samples of acute depletion of ZCCHC8 in Fig. 5 or YTHDC1 in Fig. 4, along with time-course WT (iXist-ChrX_129_, A11B2 clone) cells, cells with SPEN knockout or SPOC mutant, and cells with Xist B/C-repeat deletion (data from ref^17^). Note that X-linked gene silencing is defective in SPEN knockout, SPOC mutant, and Xist B/C-repeat deletion, thus datapoints representing those samples lie away from the predicted silencing trajectory.

## Notes

### Competing Interest Statement

The authors have declared no competing interest.

